# SoxB1 family members inhibit Wnt signaling to promote maturation and deposition of stable neuromasts by the zebrafish Posterior Lateral Line primordium

**DOI:** 10.1101/2025.04.23.650055

**Authors:** Greg Palardy, Kyeong-won Yoo, Sana Fatma, Abhishek Mukherjee, Chongmin Wang, Priyanka Ravi, Ajay B Chitnis

**Author notes:** Author for correspondence Ajay B Chitnis, Room 3B315 Building 6B 6 Center Drive, Bethesda, MD 20892.

## Abstract

Periodic formation of protoneuromasts within the migrating zebrafish Posterior Lateral Line primordium serves as a model for understanding steps that determine the self-organization of organ systems in development. Protoneuromast formation is initiated by Fgf signaling at the trailing zone of the migrating primordium in response to Fgfs produced by Wnt active cells in a leading zone. Progressive restriction of an initially broad Wnt signaling domain to a smaller leading zone allows new Fgf signaling-dependent protoneuromasts to form in the wake of the shrinking Wnt system. We show Sox2 and Sox3 are expressed in nascent and maturing protoneuromasts in a trailing part of the primordium in a pattern that is complementary to Wnt activity in a leading domain, where Sox1a, is expressed. Together, these SoxB1 factors help inhibit Wnt signaling to determine effective maturation of trailing protoneuromasts and the timely deposition of stable neuromasts. Using *dkk1b* and *atoh1b* to monitor initiation and subsequent maturation of protoneuromasts, respectively, we show how Wnt signaling regulates the pace of protoneuromast maturation in the migrating primordium, and how its inhibition by SoxB1 family members ensures maturation and deposition of stable neuromasts. Together, our observations define three steps in the periodic formation of neuromasts: first, polarization of Wnt activity in the primordium; second, pattern forming step that determines periodic formation of protoneuromasts in the context of polarized Wnt activity; and third, inhibition of Wnt signaling, which is essential for stabilizing nascent neuromasts formed in the earlier pattern forming stage.

## Introduction

The zebrafish Posterior Lateral Line (PLL) primordium is a cluster of approximately 140 cells that migrates under the skin, along the horizontal myoseptum, from near the ear to the tip of the tail, periodically forming and depositing neuromasts to spearhead formation of the zebrafish Posterior Lateral Line system (Chitnis et al., 2012). The Lateral Line is a sensory system with sensory organs called neuromasts that allow amphibians and fish to detect the pattern of water flow over their body surface. In zebrafish, nascent neuromasts, or protoneuromasts, are formed sequentially within the migrating PLL primordium starting from its trailing end. Each protoneuromast has a central hair cell progenitor, which is surrounded by cells that reorganize to form epithelial rosettes. As new protoneuromasts form progressively closer to the leading end of the primordium, previously formed protoneuromasts mature to form stable epithelial rosettes prior to deposition.

Understanding the sequence of events that coordinate specification of cell fate, shape, cellular organization and collective migration in the microcosm of the lateral line primordium contributes more broadly to understanding self-organization of tissues in a developing embryo (Ghysen and Dambly-Chaudiere, 2007). In this study we have examined the role of SoxB1 family transcription factors (Gou et al., 2018a; Gou et al., 2018b; Okuda et al., 2010; Zhang and Cui, 2014), Sox1a, Sox2 and Sox3 in the primordium, highlighting their critical role in the initiation of protoneuromast formation and maturation, enabling deposition of stable neuromasts by the migrating primordium.

Interactions between the Wnt, Fgf and Notch signaling systems (Aman and Piotrowski, 2008, 2011; Itoh and Chitnis, 2001; Lecaudey et al., 2008; Nechiporuk and Raible, 2008) coordinate periodic formation of protoneuromasts within the migrating primordium (Figure 1). The primordium forms adjacent to the ear and is dominated by Wnt/b-catenin activity, which initially extends throughout the primordium. Initiation of primordium migration toward the tail facilitates polarization of the Wnt activity (Neelathi et al., 2018) such that it becomes highest toward the leading end of the primordium (Figure 1A). Wnt signaling determines expression of factors including Wnt10 and Lef1 that locally promote Wnt activity (Aman and Piotrowski, 2008). Wnt activity also promotes expression of Fgf3 and Fgf10. However, Wnt active cells producing Fgf ligands cannot respond to them as Wnt activity promotes expression of factors that inhibit Fgf receptors from effectively responding (Aman and Piotrowski, 2008; Matsuda et al., 2013). Instead, these Fgfs activate Fgf receptors in a more distant, trailing part of the primordium with lower Wnt activity. Activation of Fgf signaling in the trailing domain initiates expression of the diffusible Wnt antagonist, Dkk1b. The combination of the local promotion of Wnt signaling coupled with longer range inhibition of Wnt by Fgf-dependent Dkk1b is thought to determine the self-organization of center-biased Fgf signaling centers that seed formation of a nascent neuromast or protoneuromast (Figure 1A) (Aman and Piotrowski, 2008; Dalle Nogare and Chitnis, 2020).

**Figure 1.**
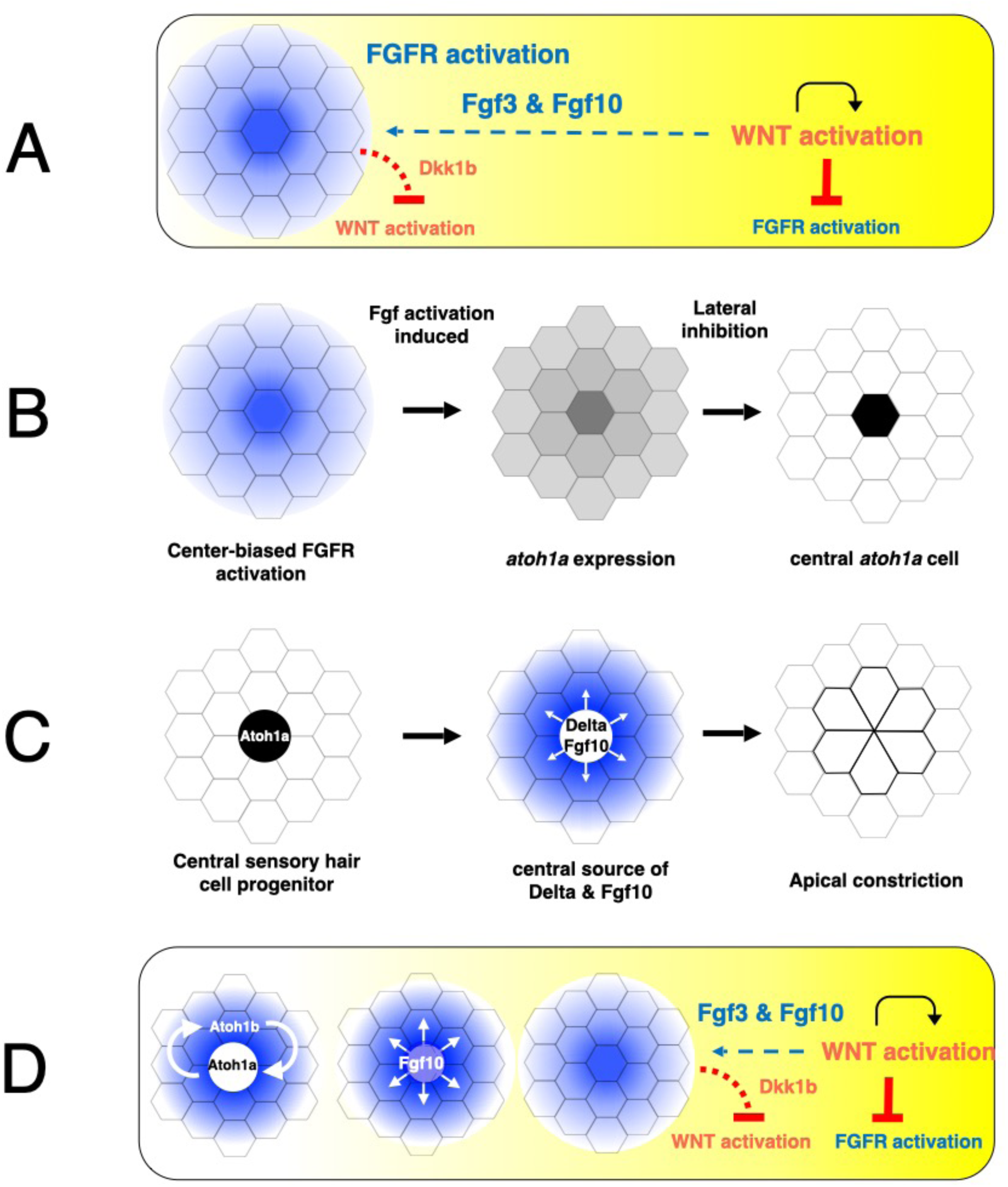
Sequence of steps involved in periodic formation of protoneuromasts. A. Interactions between Wnt and FGF signaling determine initiation of a center-biased domain of Fgf activity (blue) in the trailing part of the primordium in the context of a gradient of Wnt activity (yellow). B. Center-biased Fgf activity initiates center-biased *atoh1a* expression, which becomes restricted by Notch mediated lateral inhibition to a central cell, which becomes specified as the sensory hair cell progenitor. C. The central *atoh1a*-expressing cell becomes a source of Fgf10 and DeltaD. They activate Fgf and Notch signaling, respectively in neighboring cells to promote reorganization of neighboring cells to form epithelial rosettes. D. Maturation of the protoneuromasts is associated with establishment of *atoh1b* expression and maintenance of *atoh1a* through positive autoregulation. It is also associated with suppression of Wnt signaling in the trailing zone such that the initially broad domain of Wnt activity shrinks, creating conditions for periodic formation of new protoneuromasts in its wake. Sequential formation and maturation of protoneuromasts is associated with changes in the pattern of Fgf signaling; from center-biased in a newly formed protoneuromast, to broader as the central cell becomes a source of Fgf, to finally, donut shaped in the most mature protoneuromast, as Fgf signaling is lost in the central *atoh1a*-expressing cell.

Activation of Fgf signaling initiates the formation of a protoneuromast by coordinating both the specification of a central cell to become a sensory hair cell progenitor and the reorganization of its surrounding cells to form an epithelial rosette (Lecaudey et al., 2008; Nechiporuk and Raible, 2008). Fgf signaling initiates the expression of *atoh1a*, which gives cells in the nascent protoneuromast the potential to become sensory hair cell progenitors (Figure 1B). However, Fgf signaling and Atoh1a, respectively, also determine the expression of Notch ligands DeltaA and DeltaD (Matsuda and Chitnis, 2010). They activate Notch in their neighbors, which inhibits them from expressing *atoh1a*. As Fgf signaling and *atoh1a* is initiated in a center-biased pattern, lateral inhibition mediated by Notch signaling restricts *atoh1a* to a central cell, which becomes specified as a sensory hair cell progenitor (Chitnis et al., 2012). Activation of Fgf signaling also promotes epithelialization of primordium cells; they become taller and acquire apical basal polarity. Fgf-dependent expression of factors, like *shroom3,* promote apical constriction and the reorganization of epithelialized primordium cells to form epithelial rosettes (Figure 1C) (Ernst et al., 2012; Lecaudey et al., 2008; Nechiporuk and Raible, 2008).

The maturation of protoneuromasts at the trailing end is accompanied by progressive restriction of the initially broad Wnt domain to a smaller leading domain. As the Wnt system shrinks, another Fgf signaling domain is established in its wake, seeding formation of another protoneuromast (Figure 1D). At the same time, the central Atoh1a-expressing sensory hair cell progenitor becomes a new source of Fgf10. It activates Fgf receptors in surrounding cells, helping to consolidate their reorganization to form epithelial rosettes. The central Atoh1a-expressing sensory hair cell progenitor also expresses DeltaD (Matsuda and Chitnis, 2010), which by activating Notch in neighboring cells, promotes expression of adhesion molecules and factors that promote stability of apical cell junctions (Kozlovskaja-Gumbriene et al., 2017). In this manner, by determining expression of both Fgf and Notch ligands, the central Atoh1a-expressing sensory hair cell progenitor plays a critical role in determining maturation of stable epithelial rosettes prior to their deposition by the migrating primordium (Figure 1C).

As maturing protoneuromasts in the trailing domain become progressively separated from leading Wnt-active cells that are the initial source of Fgf signals, maintenance of Atoh1a expression must also become independent of the Fgf signals that initiated its expression in newly forming protoneuromasts. This is achieved by Atoh1a determining expression of *atoh1b*, which in turn determines *atoh1a* expression, creating a positive-feedback loop that maintains Atoh1a expression in the maturing sensory hair cell progenitor independent of Fgf signaling (Figure 1D).

Initiation and subsequent maturation of protoneuromasts within the primordium is expected to be associated with systematic changes in the pattern of Fgf signaling, where center-biased Fgf signaling initiates protoneuromast formation and *atoh1a* expression. However, as the central *atoh1a*-expressing cell becomes a source of Fgf signals, it activates Fgf signaling in its neighbors, and the pattern of Fgf signaling broadens to include neighboring cells. Eventually, as Atoh1a inhibits Fgf receptor expression, Fgf signaling is lost in within the central hair cell progenitor (Matsuda and Chitnis 2010) and the pattern of Fgf signaling is expected to become donut shaped (Figure 1D, Fgf activity in blue).

This study began with the observation that *sox2* is expressed in a changing pattern consistent with the predicted dynamic pattern of Fgf signaling in the primordium described above. However, in contrast to expectations, it was found that Fgf signaling is not directly responsible for determining sox2 expression; rather, it is the absence of Wnt signaling that facilitates *sox2* expression in the primordium. Further investigation of *sox2* function in the migrating primordium revealed its critical role in inhibiting Wnt signaling and limiting it to a leading zone in a partially redundant manner with Sox1a and Sox3. Furthermore, these SoxB1 transcription factors were shown to be essential for effective maturation of proneuromasts and timely deposition of neuromasts by the migrating primordium.

## Results

### Differential regulation of sox1a, sox2 and sox3 by Wnt and Fgf signaling in the lateral line primordium

There are three members of SoxB1 family of transcription factors expressed in the migrating primordium: *sox1a*, *sox2* and *sox3*. *sox1a* is expressed in a leading zone of the primordium (Figure 2A), where protoneuromasts are absent or just beginning to form. Conversely, *sox2* and *sox3* are most prominent in a trailing part of the primordium, which is associated with domains where protoneuromasts periodically form and mature (Figure 2B, C). *sox1a* is expressed in a relatively broad leading zone in a pattern that is like the Wnt-dependent expression of *lef1* (Figure 2D), suggesting its expression might be regulated by Wnt signaling. *sox2* and *sox3* expression in the trailing part resembles that of Fgf-dependent *pea3* (Figure 2B, C and F), consistent with their association with Fgf-dependent formation of protoneuromasts. *sox2* expression is initiated toward the leading end in a pattern consistent with its dominance at the center of the most recently formed protoneuromasts (Figure 2B, white arrows). Expression becomes broader and more intense in a more trailing domain, in association with a progressively maturing protoneuromast (Figure 2B, blue arrow). Finally, sox2 expression is often absent in the center of the most mature protoneuromast in the trailing domain of a migrating primordium (Figure 2B, black arrow). *sox3* is expressed in a pattern that is like that of *sox2* in the trailing part of the primordium, however, its dynamic changes in expression are not as easy to see as with *sox2* probes. Changes in *sox2* expression, described above, from the leading to the trailing end of the primordium are consistent with expected pattern of Fgf signaling activity in the primordium described earlier (Figure 1D). Overlap of *sox1a* expression with *lef1* and the complementary pattern of *sox2* and *lef1* expression is demonstrated with Hybridization Chain Reaction (Supplementary Figure 1).

**Figure. 2.**
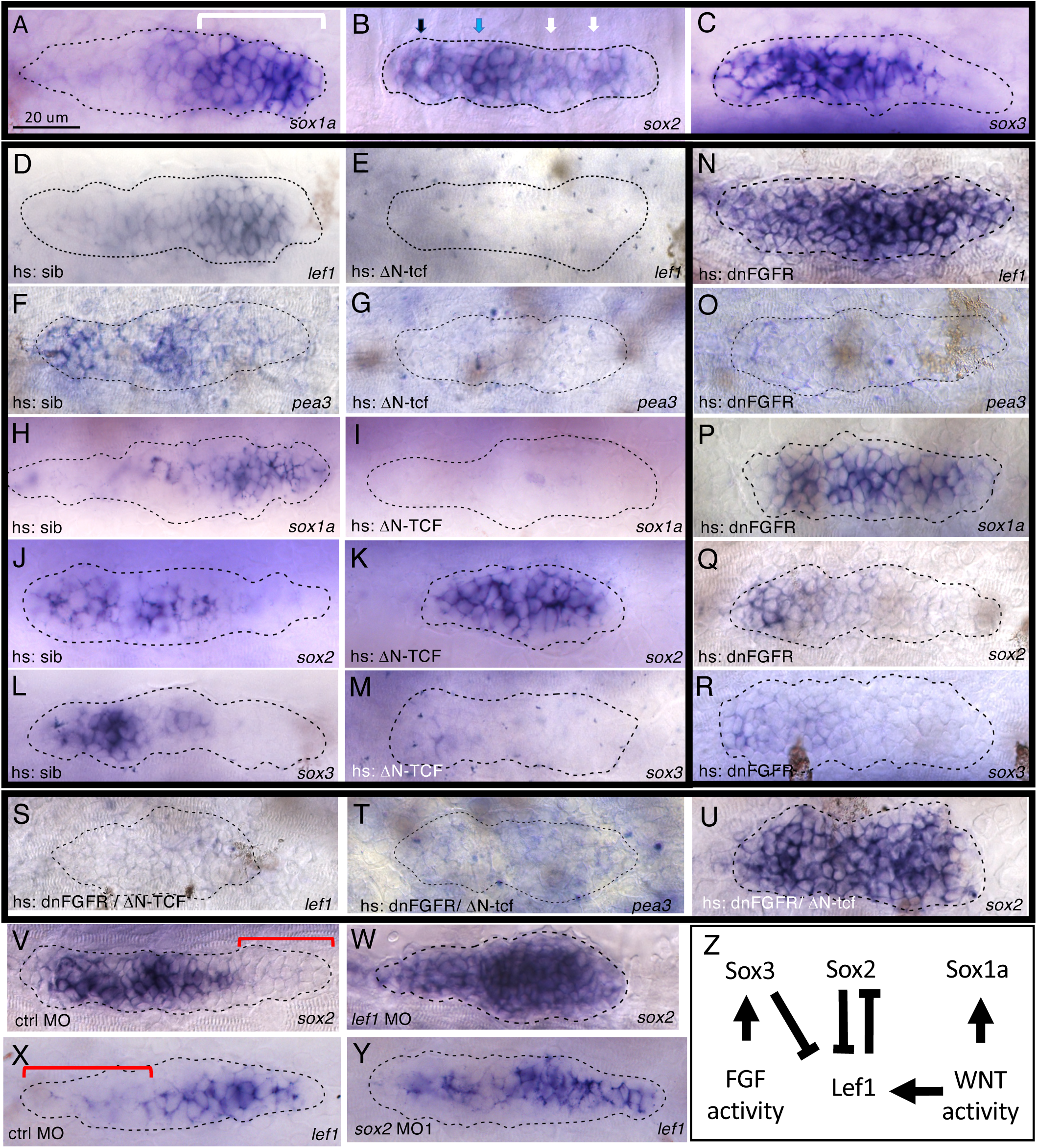
Regulation of soxB1 family members by Wnt and Fgf activity in the pLL primordium A-C. Expression of soxB1 family members in the pLL primordium of claudinB-GFP transgenic embryos. *sox1a* in the leading domain (bracket)(A), *sox2* and *sox3* in the trailing domain (B, C). Arrows in B indicate center-biased (white), broad (blue) and donut-shaped (black) sox2 expression domains. **D-M**. Effects of Heat shock induced ΔN-TCF. Expression of Wnt-responsive *lef1* and Fgf-responsive *pea3* following heat shock in wt siblings (D, F) and hs: ΔN-TCF (E, G) embryos. Expression of *sox1a*, *sox2* and *sox3* following heat shock in wt siblings (H, J, L) and hs: ΔN-TCF (I, K, M) embryos. **N-R.** Effects of heat shock induced dnFGFR expression. Expression of *lef1* and *pea3* following heat shock in hs: dn-FGFR embryos (N,O). Expression of *sox1a*, *sox2* and *sox3* following heat shock in hs: dn-FGFR embryos (P-R). **S-U.** Expression of *lef1*, *pea3* and *sox2* in following heat shock in hs: ΔN-TCF/ hs: dn-FGFR double transgenic embryos. **V, W**. *sox2* expression in control and *lef1* morphant claudinB-GFP embryos. Bracket indicates leading domain. **X, Y.** *lef1* expression in control and sox2/3 morphant claudinB-GFP embryos. Bracket indicates trailing domain. Z. Regulation of soxB1 family member expression. All embryos were fixed at 32hpf. Heat shock performed at 26-28 hpf after pLLP had clearly separated from the otic vesicle. Dotted lines demarcate outer edge of pLLP as determined by immunofluorescence(anti-GFP) or DAPI staining.

As s*ox1a* expression in the leading domain of the primordium is consistent with its expression being determined by Wnt/b-catenin signaling, and *sox2* and *sox3* expression reflects the predicted pattern of Fgf signaling, we asked if Wnt and Fgf signaling, respectively, regulate their expression. Wnt target genes can be repressed by ΔN-TCF, a dominant repressor form of TCF3 (Lewis et al., 2004) that lacks its N-terminal domain required to bind its co-activator b-catenin but retains its domain for binding a co-repressor Groucho. Heat shock induced expression of ΔN-TCF results in reduction or loss of both *lef1* and *sox1a* expression in the primordium (Figure 2D, E, H, I), supporting a role for Wnt/b-catenin signaling in determining *sox1a* expression.

Fgf signaling in the primordium is initiated by Wnt/b-catenin signaling-dependent Fgf expression by cells in the leading domain. Therefore, heat shock induced ΔN-TCF expression also inhibits Fgf expression in the primordium, which is reflected by reduced expression of Fgf-dependent *pea3* (Figure 2F, G). Though both *sox2* and *sox3* are expressed in similar patterns in the primordium, heat shock induced expression of ΔN-TCF had opposite effects on expression of *sox2* and *sox3*; *sox2* expression persists and expands into the leading zone, filling the entire primordium (Figure 2J, K), while *sox3* expression is reduced or eliminated (Figure 2L, M). The opposite effects of ΔN-TCF on *sox2* and *sox3* can be interpreted in the context of the inhibitory effects of ΔN-TCF on both Wnt and Fgf signaling. As ΔN-TCF inhibits Wnt/b-catenin, aberrant expansion of *sox2* into the leading domain suggests that Wnt/b-catenin signaling normally inhibits its expression in the leading zone. Furthermore, persistence of *sox2* expression suggests that Fgf signaling is not essential for its expression in the trailing domain, rather it is the absence Wnt/b-catenin signaling in the trailing domain that, at least in part, allows its expression there. On the other hand, loss of both *pea3* and *sox3* expression from the trailing domain following heat shock induced ΔN-TCF expression suggests that Fgf signaling is required for *sox3* expression in the trailing domain.

To further examine the role of Fgf signaling in *sox1a*, *sox2* and *sox3* expression in the primordium, we examined their expression following heat shock-induced expression of a dominant negative Fgf receptor (DNFgfR) (Lee et al., 2005). Heat shock induced expression of DNFgfR has complementary effects on the pattern of Wnt and Fgf signaling in the primordium; while it results in loss of Fgf-dependent *pea3* from the trailing domain (Figure 2O), loss of Fgf-dependent inhibition allows aberrant expansion of Wnt/b-catenin activity into the trailing domain, as reflected by expanded *lef1* expression (Figure 2N). Consistent with *sox1a* expression being determined by Wnt signaling, *sox1a* expression also expands to cover the entire length of the primordium following heat shock induction of DNFgfR (Figure 2P). On the other hand, expression of both *sox2* and *sox3* is reduced or lost in the primordium (Figure 2Q, R). However, the preceding analysis suggested their expression is lost for different reasons. While loss of Fgf signaling is likely to account for loss of *sox3* expression, the loss of *sox2* expression is likely to be related to the expansion of Wnt signaling activity into the trailing domain, where it prevents *sox2* expression. To test this interpretation, expression of *sox2* was re-examined following heat shock induced expression of both DNFgfR and ΔN-TCF, which is expected to result in simultaneous loss of both Wnt and Fgf signaling. When double transgenic HS: DNFgfR and HS: ΔN-TCF embryos were examined at 32 hpf following heat shock at 28 hpf, expression of both *lef1* and *pea3* was lost, confirming simultaneous loss of both Wnt and Fgf signaling in the primordium (Figure 2S, T). At the same time, expression of *sox2* recovered and expanded in embryos with heat shock induced expression of both DNFgfR and ΔN-TCF Wnt (Figure 2U), confirming that *sox2* expression is determined, at least in part, by the absence of Wnt signaling, rather than presence of Fgf signaling. As *sox2* and *sox3* expression is complementary to *lef1*, we asked if they mutually inhibit each other. Consistent with this, knockdown of *lef1* allowed *sox2* expression to expand into the leading zone, while knockdown of *sox2* and *sox3* with sox2MO1 (described below) allowed *lef1* to expand into the trailing zone (Figure 2V-Y). In summary, while expression of *sox1a* is determined by Wnt activity, expression of *sox3* is determined by Fgf signaling, and *lef1* and *sox2* mutually inhibit each other’s expression (Figure 2Z)

### Sox2 is required for timely deposition of the first stable neuromast

A previously published morpholino (sox2MO1: GCTCGGTTTCCATCATGTTATACAT), designed to inhibit translation of *sox2* (Ogai et al., 2014), was initially used to assess function of Sox2. Knock-down of *sox2* resulted in absence or delayed deposition of the first and sometimes second and third neuromasts (Figure 3A, B, E, F). Furthermore, the primordium deposited fewer neuromasts by the time its migration ended at the tip of the tail with the formation of terminal neuromasts. The initial observations made in *sox2* morphants were confirmed with CRISPR generated sox2^y589^ mutants (Figure 3C, D, G, H) in which a 14 base pair insertion resulted in the introduction of a premature stop codon in the first exon, eliminating effective translation of its functional domains including the High Mobility Group and Transactivation domain (Supplementary Figure 4A).

**Figure 3.**
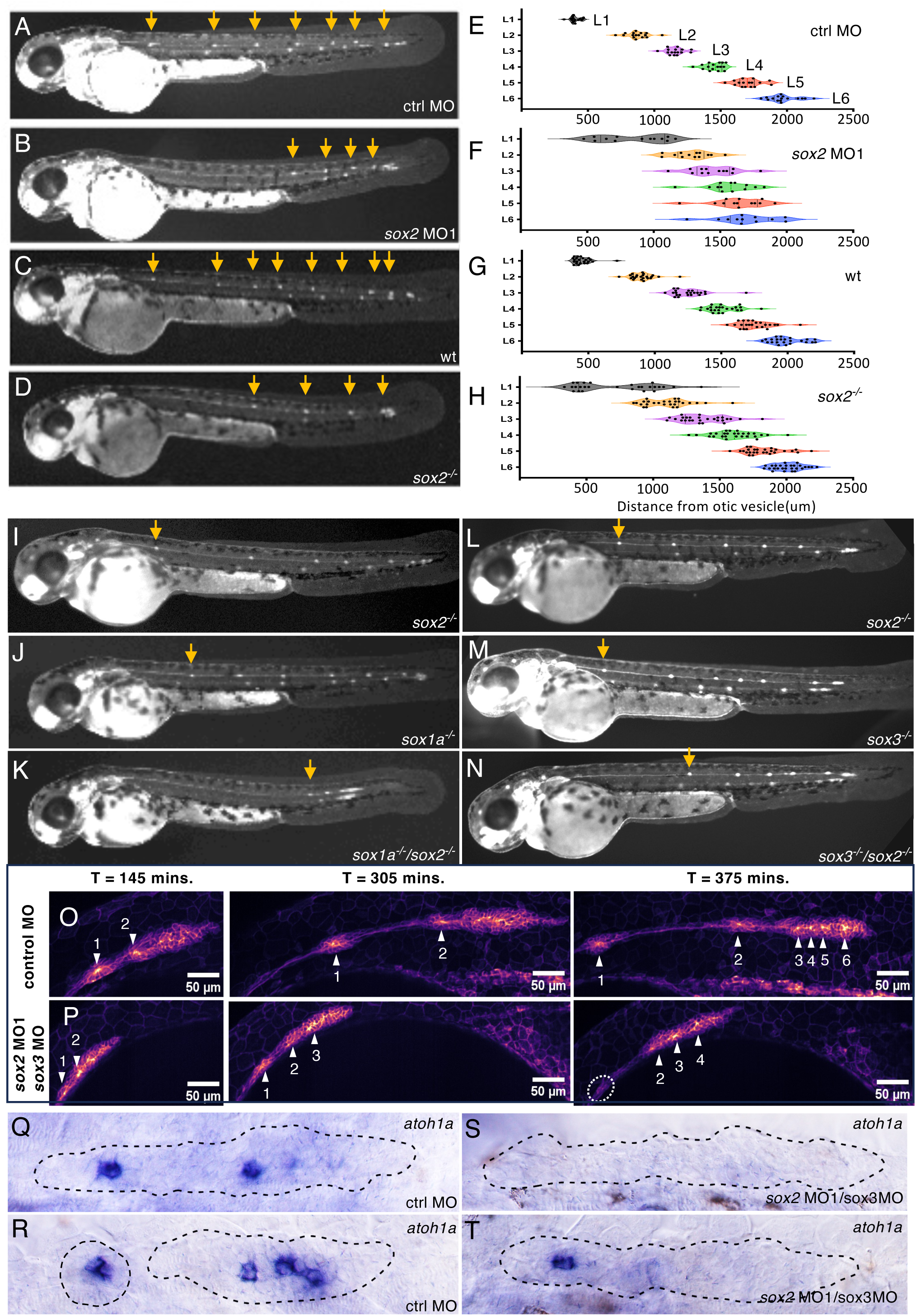
Deposition pattern of neuromasts in soxB1 family morphants and mutants in claudinB-GFP embryos A-D. Deposition pattern of neuromasts (arrows): in a control morpholino (ctrl MO) and 2ng sox2 MO1 morpholino injected embryo (A,B), and in wild type sibling (wt) and sox2^-/-^ mutant (C,D). **E-H.** Distances of L1-L6 neuromasts from the otic vesicle in embryos represented in A-D, respectively. **I, J, K.** Neuromast deposition pattern in a sox2^-/-^, sox1a^-/-^, and sox1a^-/-^/sox2^-/-^ double mutant embryo. **L, M, N** Neuromast deposition pattern in a sox2^-/-^, sox3a^-/-^, and sox2^-/-^/sox3^-/-^ double mutant. Arrows in I-N indicate position of 1^st^ neuromast (L1). All imaged at 52 hpf. **O, P.** Live imaging of early deposition events in control and sox2MO1/sox3 MO injected embryos. Numbered Arrowheads indicate position of deposited neuromast or forming protoneuromasts. Dotted oval outlines disassembled deposited neuromast 375 minutes after initiation of image acquisition at 25hpf. **Q-T.** Expression of *atoh1a* in control and sox2MO1/ sox3 MO co-injected embryos at 26hpf.

Though *sox2* mutants initially recapitulated observations in the sox2MO1 morphants (Figure 3B, D), after maintenance of the mutant for several generations, the severity of the phenotype reduced and was not fully penetrant in homozygous sox2^y589^ mutant embryos (compare Figure 3D, I, L). This raised the possibility that other SoxB1 family members expressed in the primordium had the potential to partially compensate for loss of *sox2* in the primordium. To examine the role of *sox1a* and *sox3* in potentially compensating for loss of *sox2*, a previously generated *sox3* mutant was acquired (Gou et al., 2018a) and a CRISPR-generated *sox1a^y590^*mutant was generated in which a 13 base pair deletion introduces a stop codon in the first exon (Supplementary Figure 4B). Comparing the pattern of neuromast deposition in homozygous *sox2* mutants with *sox2, sox1a* double or *sox2*, *sox3* double homozygous mutants showed that simultaneous loss of *sox2* with either *sox1a* or *sox3* results in a more severe phenotype characterized by a greater fraction of embryos showing delayed deposition of the first neuromast (Figure 3K, N). It should be noted that as the original sox2^y589^ mutant line has been maintained for multiple generations, the homozygous mutant phenotype has become progressively weaker and only a small fraction of homozygous mutant embryos now show delayed deposition of L1, as opposed to roughly half when the y589 line was first created (compare position of L1 in sox2^-/-^ in Figure 3D and Figure 3I or L). Keeping this in mind, neither homozygous *sox1a*, *sox2*, or *sox3* mutants on their own had a significant fraction of embryos with delayed deposition of L1. On the other hand, simultaneous loss of *sox2* and *sox1a* function resulted in a much more severe delay in deposition of the first neuromast and at approximately 48-52hpf the first obvious neuromast was typically observed at a much more caudal position (Figure 3K). *sox2/sox3* double mutants, on the other hand, had a phenotype like that seen in *sox2* MO1 morphants (Figure 3B, N).

Though the similarity of *sox2/sox3* double mutants to the phenotype seen in sox2MO1 morphants was initially attributed to potential differences in genetic compensatory mechanisms operating in morphants versus mutants, our analysis of the previously published sox2MO1 morpholino revealed that its sequence allowed it to simultaneously target *sox3*. Comparison of effects of sox2MO1 and *sox3* MO on Sox2 and Sox3 expression with antibody staining confirmed that the sox2MO1 effectively inhibited both Sox2 and Sox3 expression, while the sox3 MO selectively inhibited Sox3 expression without any detectable effect on Sox2 expression (Supplementary Figure 2A). An alternate sox2 morpholino (AACCGATTTTCTCGAAAGTCTACCC), shown to be more specific for Sox2 (Kamachi et al., 2008), had a less penetrant effect on the pattern of L1 deposition (Supplementary Figure 2B). This confirms that Sox2 and Sox3 partially compensate for each other’s function and that the effects of *sox2* knockdown alone are not fully penetrant. As the specific *sox2* morpholino had mild effects and would have to be co-injected with an additional *sox3* specific morpholino to observe the more penetrant phenotype, this study continued use of the dual effect sox2MO1 to simultaneously knockdown *sox2* and *sox3* function. As some of the studies were performed before it was realized that sox2MO1 simultaneously reduces *sox2* and *sox3* function, data that includes co-injection of sox2MO1 and sox3 MO has been included in this study.

Analysis of the pattern of neuromast deposition in embryos with homozygous and heterozygous wild-type and mutant genotypes provided further evidence of potential interactions between these soxB1 family members. For example, while *sox1a^-/-^, sox2^+/+^* embryos had no significant delay in deposition of L1, *sox1a^-/-^, sox2^-/+^* mutants had many more embryos with a delay (Supplementary Figure 3A). Similarly, while neither *sox2^-/+^* or *sox3^-/+^* heterozygous mutants embryos showed delays in an otherwise wild-type background, delays were observed in *sox2^-/+^*, sox3^-/-^ or *sox2^-/-^, sox3^-/+^* embryos (Supplementary Figure 3B).

Live imaging of the *sox2MO1*/*sox3* double morphants showed that the morphant primordium migrates slower and L1 deposition was delayed in 10/10 cases (Figure 3O, P). Furthermore, in 6/10 cases, initial neuromasts were poorly formed, unstable and fell apart (Figure 3O, P, movies 1-3). In 5/6 of these, an unstable poorly formed rosette was initially deposited but some of its cells reassociated with the primordium to continue their migration. This suggests that in the morphants, a neuromast may have formed and deposited but was either dissolved or its cells reassociated with the migrating primordium, preventing observation at 48-52 hpf when the pattern of deposited neuromasts are typically examined. Hence many neuromasts labelled L1 in morphants may correspond to later deposited neuromasts that were more stable than earlier deposited ones.

Individual protoneuromasts, as well as recently deposited neuromasts, are each marked by a central *atoh1a-*expressing cell. Examination of *atoh1a* showed that delay in deposition of the first neuromast was also related to fewer *atoh1a*-expressing cells within the migrating primordium. At 26hpf, 9/9 control primordia were associated with at least two *atoh1a*-expressing cells (Figure 3Q, R). Of these, 4/9 had two prominent *atoh1a* cells, either both within the migrating primordium (Figure 3Q), or with one in the trailing domain and other in a recently deposited neuromast. The remaining 5/9 primordia were associated with four *atoh1a*-expressing cells, either all within the primordium or in the primordium and in a recently deposited neuromast (Figure 3R). In *sox2MO1/sox3* double morphants on the other hand, 3/13 primordia were associated with just one trailing *atoh1a* expressing cell, two of which were within depositing neuromasts (Figure 3T). The remaining 10/13 primordia had no *atoh1a*-expressing cell and no deposited neuromasts (Figure 3S). Together these observations show that combinatorial loss of *sox2* and *sox3* results in both a delay in sequential formation of protoneuromasts within the migrating primordium and an instability of neuromasts that are deposited.

### Loss of Sox2 with Sox1a or Sox3 results in expansion of Wnt activity, not a loss of Fgf signaling

As protoneuromast formation is dependent on suppression of Wnt and establishment of stable Fgf signaling centers in the wake of the shrinking Wnt system, the expression of *lef1* and *pea3* was examined to determine if changes in Wnt and Fgf signaling systems, respectively, contribute to the delay in the stable formation and deposition of the first neuromast. Knock down of *sox2* and sox3 with sox2MO1 resulted in expansion of Wnt activity to the trailing domain (Figure 4A, B). Loss of either *sox1a*, *sox2* or *sox3* function on their own had little effect on *lef1* expression. However, loss of *sox2* in combination with either *sox1a* or *sox3* led to expansion of *lef1* into the trailing zone. Nevertheless, the expansion of *lef1* was not accompanied by a loss of *pea3* expression, suggesting that the delay in deposition of a stable neuromast primarily correlates with expanded Wnt signaling, not the loss of Fgf signaling (Figure 4C-N). Though potential expansion of Wnt activity as evidenced by expansion of *lef1* expression was not seen in sox2^-/-^ primordia, *sox1a* expanded into the trailing zone (Figure 4O, P). This suggested that Wnt activity had indeed expanded following loss of *sox2* alone and that *sox1a*, whose expression is also dependent on Wnt activity, may serve as a more sensitive indicator of its activity.

**Figure 4.**
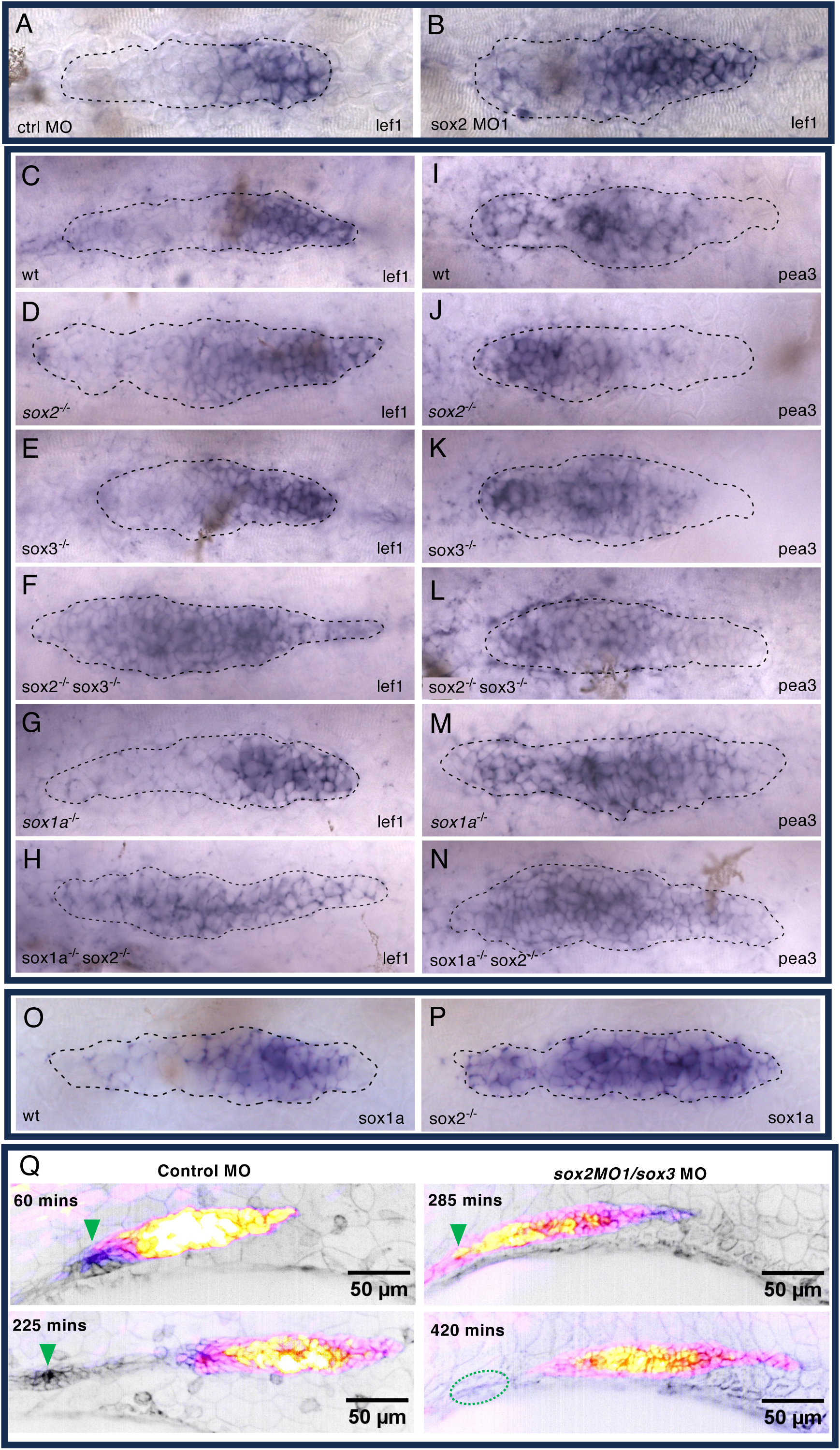
soxB1 family members inhibit Wnt activity in the pLL primordium A, B. Expression of *lef1* in control and sox2 MO1 morpholino injected embryos. **C-H.** Expression of *lef1* in soxB1 family single and double mutants. **I-N.** Expression of *pea3* in soxB1 family single and double mutants. **O, P.** Expression of *sox1a* in wt and *sox2* mutant embryos. Dotted lines demarcate outer edge of pLLP. **Q.** Time-lapse images of Wnt activity in control and sox2/sox3 morphants in *Tg[tcf/lef1-miniP:d2GFP] / CldnB:Lyn-mScarlet* transgenic embryos. GFP reporting Wnt activity visualized with Fire LUT (Yellow to white-high, purple -low) and cell membranes with Lyn-mScarlet (inverted grey scale). Arrowhead indicates depositing neuromast. Dotted oval outlines disassembled deposited neuromast 420 minutes after initiation of image acquisition at 25hpf.

The expansion of Wnt activity in the primordium in embryos injected with *sox2* morpholinos in combination with either *sox1a* or *sox3* morpholinos was also examined in live embryos by examining changes in GFP expression in the Tcf/Lef-miniP:d2eGFP Wnt reporter line (Shimizu et al., 2012). As seen previously with live imaging, co-injection of *sox2* and *sox1a* or *sox3* morpholinos resulted in formation of unstable neuromasts that typically fell apart following deposition. However, in this context it was possible to see that instability of neuromasts was associated with expanded Wnt-activity-dependent GFP expression that extended to the trailing end of the migrating primordium and, at least transiently, in cells of the depositing neuromasts (Figure 4Q, movie 4). The time lapse also showed that primordium migration was initiated late and was slower in the morphants. As a result, images of the primordia at similar positions in their migration above the yolk sac in Figure 4Q were captured at different times, 60 and 225 minutes for controls, and 285 and 420 minutes in morphants.

### Inhibition of Wnt activity restores more timely deposition of the first neuromast

As the delay in deposition of neuromasts appeared to be associated with expanded Wnt activity, we asked if inhibiting Wnt activity would help restore timely deposition of L1. When embryos injected with either *sox2* MO1 morpholinos alone or in combination with *sox3* morpholinos were compared with similar embryos exposed to 30 uM IWR (Chen et al., 2009) between 28 and 32 hours, expansion of *lef1* expression was reduced (Figure 5A-D). Similarly, delays in deposition of the L1 neuromast were found to be significantly reduced by exposure to IWR (Figure 5E-I).

**Figure 5.**
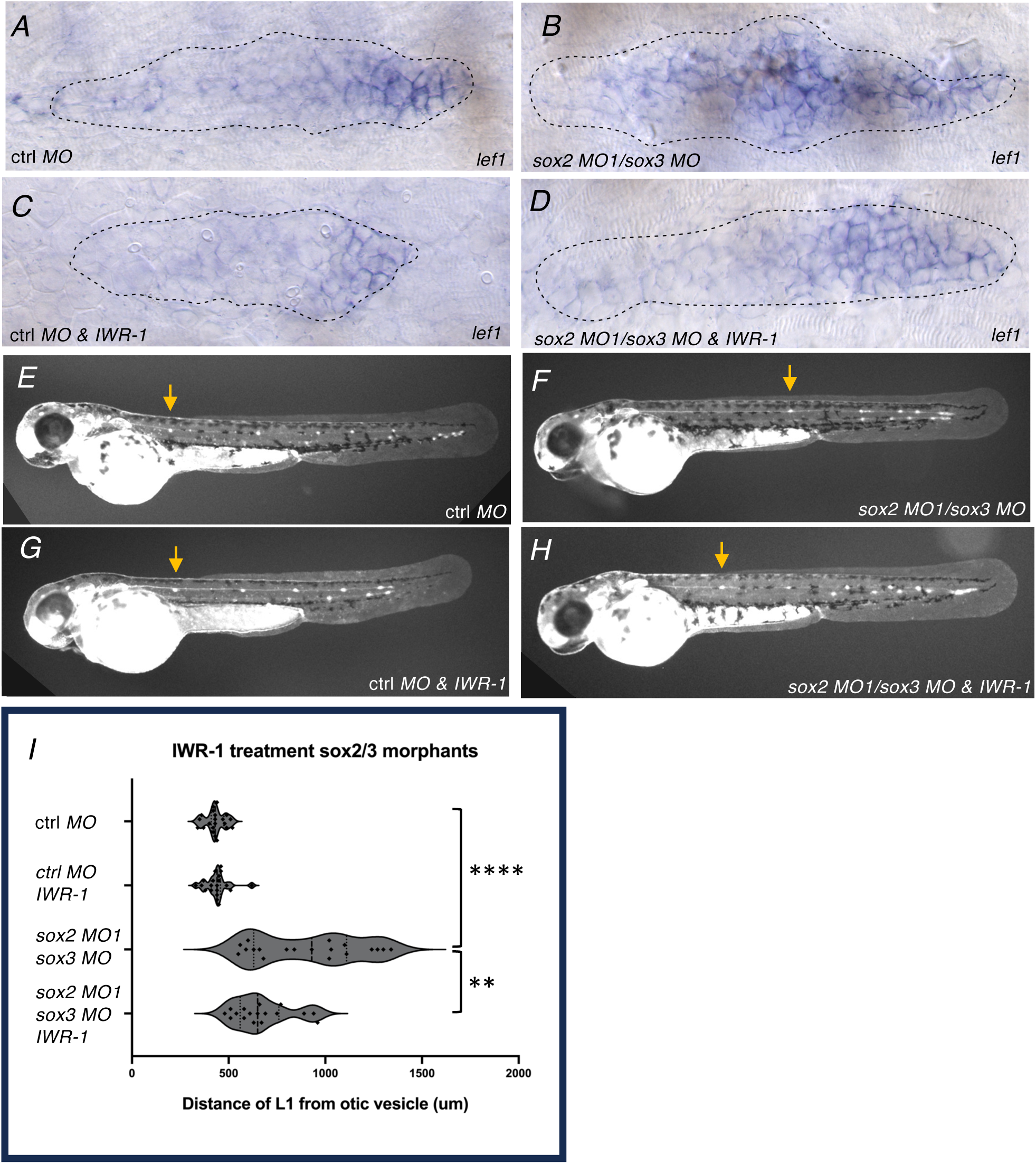
Inhibition of Wnt activity in sox2/sox3 morphants restores timely L1 deposition A-D. Expression of *lef1* in control and sox2 MO1-injected embryos treated with DMSO (A, B) or Wnt signaling antagonist IWR-1(C, D). **E-H.** Deposition pattern of neuromasts for same treatment groups. Arrow indicates position of 1^st^ deposited neuromast L1. I. Quantification of distance of L1 from otic vesicle for treatment groups. Asterisks indicate significance by Mann-Whitney test (**** p< 0.0001, ** p= 0.0079).

### Loss of Sox2 with Sox1a or Sox3 prevents effective maturation of protoneuromasts

As nascent rosettes mature in the trailing part of the primordium, Atoh1a initiates expression of *atoh1b* and begins to sustain its own expression through autoregulation. At the same time, in the trailing part of the primordium, there is a progressive loss and eventual absence of *dkk1b* expression in the maturing protoneuromast. Consequently, the sequential formation and maturation of protoneuromasts can be visualized by simultaneously examining expression of *dkk1b*, whose expression marks nascent proneuromasts, while its disappearance, coupled with appearance of *atoh1b*, labels maturing protoneuromasts (Figure 6A).

**Figure 6.**
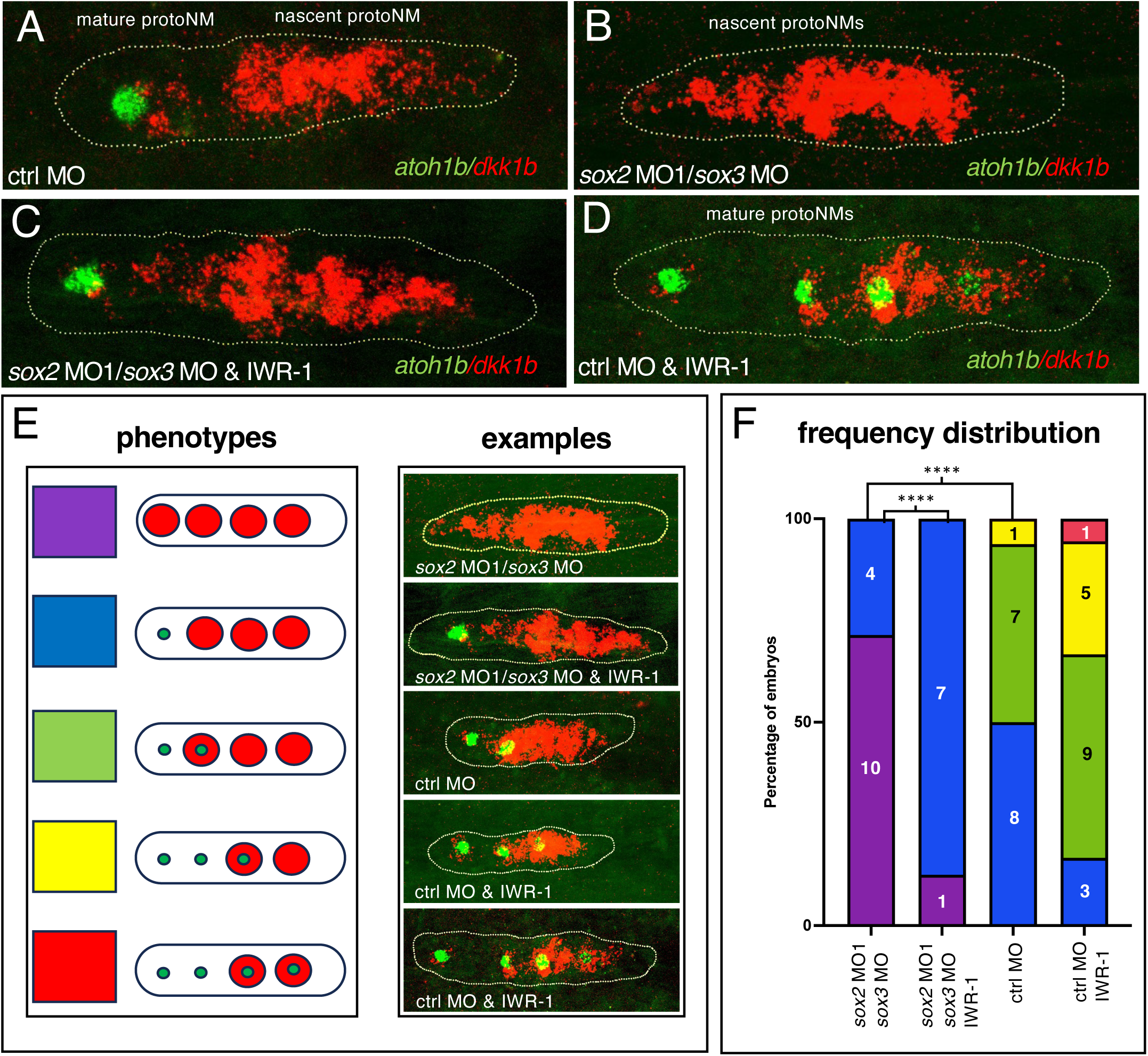
Inhibition of Wnt activity in sox2/3 morphants restores timely neuromast maturation A-D. Expression of *dkk1b* and *atoh1b*, which, respectively, mark nascent and mature protoneuromast, in control and sox2 MO1morphant embryos, in the absence or the presence of the Wnt signaling antagonist IWR-1. **E.** Schematized phenotypes with corresponding examples of patterns of *dkk1b* (large red circle) and *atoh1b* (small green dot) expression. Each pattern is coded by a color. F. Frequency distribution of phenotypic categories in treatment groups using the color code. Asterisks indicate significance by Fisher’s exact test (**** p< 0.0001)

Knockdown of *sox2* function together with *sox1a* (data not shown) or *sox3*, following morpholino injections resulted in loss of *atoh1b* expression and expansion of *dkk1b* expression to more trailing parts of the primordium (Figure 6B-F). Together with previous results, this observation suggested that expansion of Wnt activity into the trailing zone, following knockdown of *sox2* and *sox1a* or *sox3*, was preventing effective initiation of *atoh1b* expression and allowed *dkk1b* expression to persist in the trailing zone. To test this hypothesis, we asked if suppression of Wnt activity, which restores more timely L1 deposition, also allows more effective maturation of trailing protoneuromasts. Knockdown of *sox2* together with *sox3*, coupled with exposure to IWR, increased the fraction of primordia with trailing *atoh1b* cells and less *dkk1b* in the trailing zone (Figure 6C-E). Furthermore, exposure to IWR in control embryos shrunk the *dkk1b* domain to a smaller leading domain and increased the number of protoneuromasts associated with *atoh1b* expression (Figure 6D-F). It also increased the number of transitory protoneuromasts associated with both *atoh1b* and *dkk1b* expression. Together these observations suggest that *sox2* functions in a partially redundant role with *sox1a* and *sox3* to inhibit Wnt activity in the trailing part of the primordium. Thus, SoxB1-dependent inhibition of Wnt activity sets the pace for effective maturation of nascent protoneuromasts and their deposition as stable epithelial rosettes.

## DISCUSSION

This study has examined the expression and function of three members of the SoxB1 family of transcription factors, Sox1a, Sox2 and Sox3, in the migrating posterior lateral line primordium. The expression of each of these SoxB1 family members is regulated in a distinct manner by Wnt and FGF signaling. We discuss how progressive changes in their expression along the length of the migrating primordium reflects the dynamics of Wnt and FGF signaling in primordium, shedding light on the way interactions between these signaling pathways initiate formation and subsequent maturation of protoneuromasts within the migrating primordium. Functional analysis revealed how these SoxB1 transcription factors operate as part of a robust regulatory network to limit Wnt activity, ensuring progressive shrinking of the Wnt system and the timely formation and maturation of sequentially formed neuromasts prior to their deposition. Finally, we discuss how previous studies describing the establishment of polarized Wnt activity and self-organization of protoneuromast in the context of this gradient, together with this study, which describes the collective role played by Sox1a, Sox2 and Sox3 in determining maturation and deposition of stable neuromasts, helps define more broadly key steps that contribute to self-organization of organ systems during animal development.

### *Sox1a*, sox2 and *sox3* are expressed and regulated in distinct and complementary ways by Wnt and Fgf signaling

*sox1a* expression is determined by Wnt signaling in a leading part of the primordium, while expression of *sox2* and *sox3* is restricted to the trailing zone in association with the initial formation and subsequent maturation of protoneuromasts. Though *sox2* and *sox3* are expressed in similar patterns, expression of *sox3* is determined by Fgf signaling, while expression of *sox2* is determined, at least in part, by the absence of Wnt signaling and is not directly dependent on Fgf signaling. However, as the Wnt and Fgf signaling systems locally inhibit each other in the primordium, their activity patterns are for the most part complementary, though there is overlap where new protoneuromast form. Hence the pattern of *sox2*, defined by the absence of Wnt signaling, is complementary to that of *sox1a* and like that of *sox3*, whose expression is determined more directly by Fgf signaling. It remains unclear what more directly promotes *sox2* expression in the primordium.

### Sox2 expression reflects a predicted pattern of FGF and Wnt signaling in protoneuromasts

Fgf signaling is expected to have three distinct patterns in protoneuromasts from their initiation to maturation (Figure 1) (Dalle Nogare and Chitnis, 2020). Protoneuromast formation is initiated with center biased FGF signaling, as this sets the stage for initiating center-biased *atoh1a* expression, which, in turn, becomes restricted to a central cell by lateral inhibition mediated by Notch signaling. The pattern of FGF signaling would then be expected to broaden in a second phase, as protoneuromasts mature and the central *atoh1a* expressing cell becomes a source of Fgf10 that activates FGF signaling in neighboring cells. Finally, in a third phase, as the expression of Atoh1a becomes self-sustaining via autoregulation with Atoh1b, the *atoh1a*-expressing central cell stops expressing FGF receptors and Fgf signaling is expected to be lost in this central cell resulting in a donut shaped pattern of FGF activity. These predicted patterns, from center biased, to broader, and finally broader but with a central hole, are observed with *sox2* expression, even though they are determined by the absence of Wnt signaling rather than the presence of FGF signaling. Fgf-dependent *pea3* expression, often used as a proxy for the pattern of Fgf signaling, had previously been shown to be initiated in a center-biased pattern (Chitnis et al., 2012). However, this transient pattern is often difficult to see with *pea3*, and it is typically only observed in a subset of primordia. Recapitulation of the three predicted patterns of Fgf signaling with *sox2* expression lends support to a framework that describes a predicted progression of changes in the spatial pattern of Wnt and Fgf signaling from initiation of protoneuromast formation to their maturation prior to neuromast deposition (Dalle Nogare and Chitnis, 2020).

### SoxB1 family members together form a robust network to limit Wnt activity in the primordium

Loss of function studies with morpholinos and mutants showed that soxB1 transcriptions factors Sox1a, Sox2 and Sox3 work together to attenuate Wnt signaling in the primordium, guaranteeing a progressive reduction in Wnt activity from the leading to trailing end. This ensures effective suppression of Wnt signaling at the trailing end, which this study suggests is required for establishing autoregulation of Atoh1a and stabilization of epithelial rosettes, both of which are required for effective maturation of protoneuromasts prior to their deposition as neuromasts. While SoxB1 family members share a role in attenuating Wnt activity, the distinct mechanisms that determine their expression are likely to contribute to a robust network that ensures effective suppression and regulation of Wnt activity. Though *sox2* and *sox3* are expressed in a similar pattern in developing protoneuromasts, their expression, dependent on the absence of Wnt, or presence of Fgf activity, respectively, may contribute to limited function of at least one or the other when either of these regulatory mechanisms is selectively compromised. Furthermore, while *sox2* and *sox3* cooperate to inhibit Wnt activity in a trailing domain, Wnt-dependent expression of *sox1a* in the leading zone is likely to limit the level of activity that can be achieved by mechanisms that locally promote Wnt signaling. These distinct mechanisms that inhibit Wnt signaling in the leading and trailing zones appear to be synergistic, as combined interference with *sox1a* and *sox2* function has a much more severe effect on delaying stable neuromast deposition than inhibiting *sox2* and *sox3* together.

### Exaggerated Wnt activity delays protoneuromast formation and prevents effective maturation

The delay in deposition of the first stable neuromast, associated with expanded Wnt activity reflects two problems: First, it caused a delay in initiation of protoneuromast formation, as effective inhibition of Wnt signaling at the trailing end is essential to initiate Fgf-dependent protoneuromast formation, second, it prevented initiation of changes associated with maturation of protoneuromasts. The delay in initiation was evident from the absence or delay in appearance of protoneuromast-marking *atoh1a* spots in sox2/sox3 morphant primordia at 26hpf. Expanded Wnt activity also prevented initiation of changes associated with maturation of protoneuromasts. Maturing neuromasts are characterized by expression of *atoh1b* and absence of *dkk1b*, whose expression marks newly formed protoneuromasts. Both the failure to initiate *atoh1b* expression and persistence of *dkk1b* appear to be associated with expanded Wnt activity, as inhibition of Wnt activity allows reappearance of *atoh1b* and suppression of *dkk1b* in the trailing protoneuromast. The expansion of *dkk1b* expression with expanded Wnt activity is consistent with its typical expression in nascent protoneuromasts that are proximal to domains of Wnt activity where there is likely to be some overlap between Wnt and Fgf signaling activity. The failure to initiate *atoh1b* expression in *sox2/sox3* morphants suggests that Atoh1a is only able to initiate *atoh1b* expression when the Wnt activity drops below some threshold. This result also suggests that Sox2, along with other SoxB1 family members, like Sox3, have a critical role in suppressing Wnt activity below a threshold to allow initiation and effective maturation of protoneuromasts in the trailing zone.

### Three steps to self-organization of pattern and form in development

SoxB1 family members Sox1a, Sox2 and Sox2 are part of gene regulatory network that ensures effective maturation of newly formed protoneuromasts prior to their deposition as neuromasts with an organization and structural stability that sets the stage for their further growth and development. Together with previous studies, this study helps to define three steps that determine the robust and reproducible self-organization of pattern, fate and form in the migrating primordium. First, caudal migration of the primordium initiates polarization of Wnt signaling, which becomes highest at the leading end of the primordium (Neelathi et al., 2018) (Figure 7A). Second, in the context of polarized Wnt activity, local promotion of Wnt activity, coupled with its long-range inhibition by FGF signaling, determines self-organization of a center-biased domain of Fgf signaling at the trailing end of the primordium (Dalle Nogare and Chitnis, 2020) (Figure 7B). Fgf signaling initiates center-biased *atoh1a* expression, which gives cells the potential to become Hair Cell progenitors, while lateral inhibition, operating in this context, restricts *atoh1a-*dependent progenitor fate to a central cell (Chitnis et al., 2012; Lecaudey et al., 2008; Matsuda and Chitnis, 2010; Nechiporuk and Raible, 2008).

**Figure 7.**
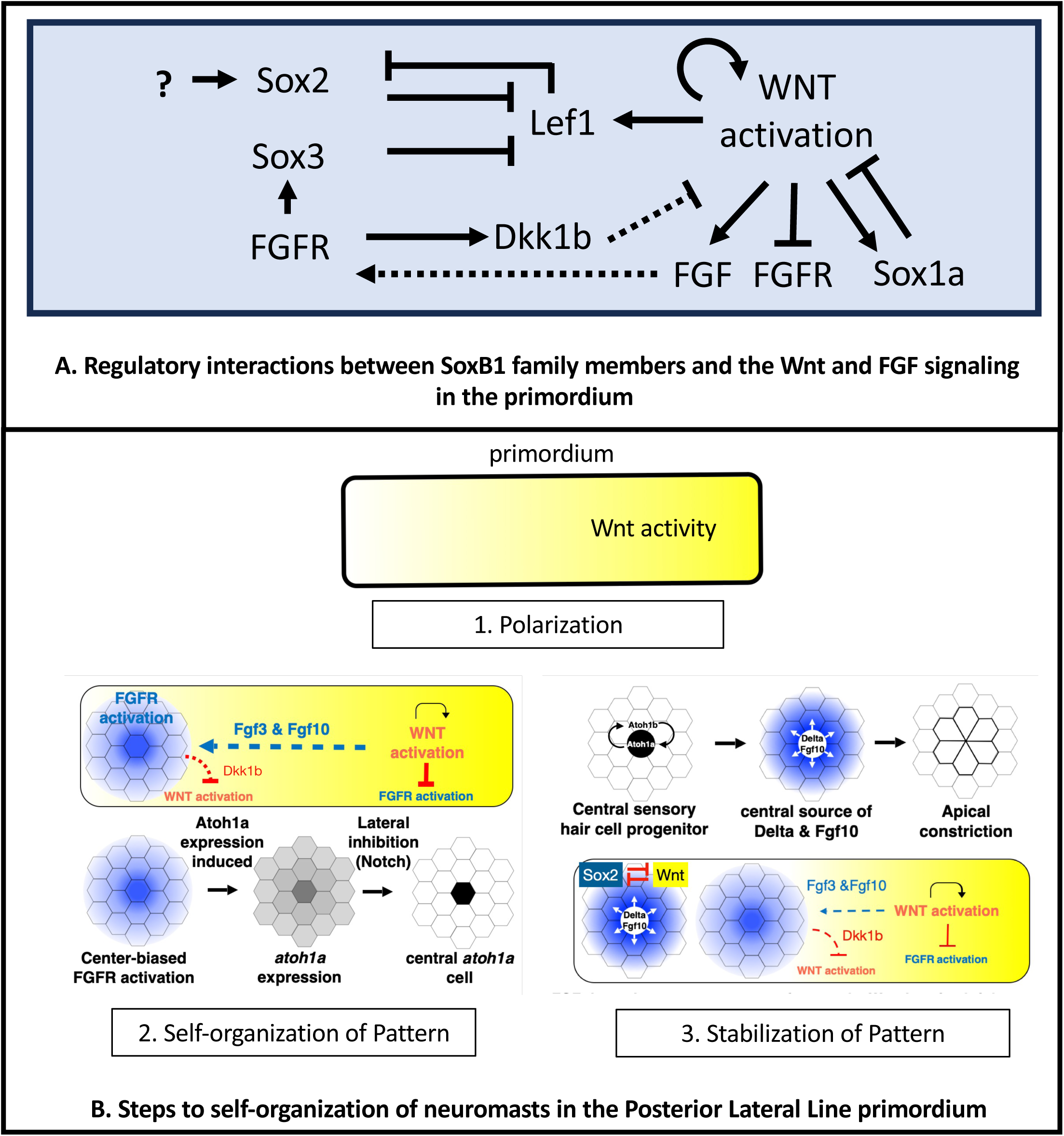
Summary of regulatory interactions and Steps to self-organization of neuromasts in the Posterior Lateral Line primordium. A. Regulatory interactions between SoxB1 family members and the Wnt and FGF signaling in the primordium B. Steps to self-organization of neuromasts in the Posterior Lateral Line primordium. See text for details.

Finally, in a third step, SoxB1 transcription factors cooperate to inhibit Wnt activity at the trailing end (Figure 7C). This allows Atoh1a to initiate *atoh1b* expression and stabilize its own expression through autoregulation. Atoh1a in turn drives FGF10 and DeltaD expression, activating Fgf and Notch signaling in surrounding cells, consolidating their reorganization to form a stable epithelial rosette (Ernst et al., 2012; Kozlovskaja-Gumbriene et al., 2017) prior to deposition. Furthermore, restricting Wnt activity to a smaller leading domain recreates the conditions for formation of another protoneuromast in the wake of the shrinking Wnt system. We suggest the three steps in the development of the primordium: first, polarization of the tissue; second, the operation of self-organizing mechanisms that determine pattern and form in the context of this polarized tissue; and finally, the stabilization of patterns and forms initiated by self-organizing mechanisms, define more broadly, a framework for understanding robust and reproducible emergence of pattern, fate, form and cell behavior in development.

## MATERIALS AND METHODS

### Fish maintenance and transgenic lines

Danio rerio were maintained under standard conditions. Embryos were staged according to Kimmel et al. 1995 (Kimmel et al., 1995). *Tg[cldnb:lynGFP]* (Haas and Gilmour, 2006), *Tg[cldnb:lyn-mscarlet]* (Dalle Nogare et al., 2020), *Tg[hs:dkk1b]* (Stoick-Cooper et al., 2007), *Tg[hsp70l:dnfgfr1-EGFP]pd1* (Lee et al., 2005), *Tg[hs:ΔNTCF:GFP]* (Lewis et al., 2004) and *Tg[tcf/lef-miniP:dGFP]* (Shimizu et al., 2012) were previously described. Heat shock was carried out at 38.5°C for 60min. Experiments were carried out in accordance with National Institute of Child Health and Human Development Animal Care and Use Committee Protocol Number 21-013.

### Morpholino (MO) injections and drug treatments

All MOs were obtained from Gene Tools LLC. MOs used were (5′-3′): *sox1a*, CCGTTTCCATCATCATGCTATACAT, (Gerber et al., 2019); sox2 MO1, GCTCGGTTTCCATCATGTTATACAT, (Ogai et al., 2014); sox2 MO2, AACCGATTTTCTCGAAAGTCTACCC, (Kamachi et al., 2008); sox2 MO3, GAGAGGCTGCTGAAGTTACCTTAGC, (Kamachi et al., 2008); sox2 MO4, GAAAGTCTACCCCACCAGCCGTAAA, (Kamachi et al., 2008); sox3, AGCTCAAACTCTGGTCAAGTAAAGT (this paper); p53, GCGCCATTGCTTTGCAAGAATTG, (Robu et al., 2007) Standard control, CCTCTTACCTCAGTTACAAATTTATA, (Gene Tools, LLC)

MOs were injected at the single-cell stage with co-injection of 1.5 ng of p53-MO. Standard control MO was injected at the same concentration as the experimental morpholino. Embryos were incubated with 10 μm SU5402 (Tocris, cat. # 3300), 5 μm BIO (Sigma cat. # 361552) or 30uM IWR-1(Sigma, cat. # I0161) from 26 hpf until fixation. Drugs were made up in DMSO, and control embryos were incubated for similar periods in DMSO alone.

### soxB1 mutant generation

*sox1a^y590^* and *sox2^y589^* mutant lines were generated by CRISPR/Cas9 targeted genome editing relying on non-homologous end joining repair mechanism, as described in detailed protocols provided by Auer et al. (Auer et al., 2014), Gagnon et al (Gagnon et al., 2014), and Talbot and Amacher (Talbot and Amacher, 2014). Optimal target sites were selected using the CHOPCHOP web tool (Montague et al., 2014). Cas9 protein with nuclear localization signal was obtained from PNABio. Guide RNAs were generated by PCR and transcribed using HiScribe T7 High-Yield RNA Synthesis Kit (NEB). 250 pg of guide RNA and 400-500 pg of *cas9* protein per embryo was injected at the one-cell stage into the cell. Putative F1 mutants were screened for indels by fluorescent PCR with fragment analysis on an ABI3730XL (Carrington et al., 2015) and later confirmed by Sanger sequencing. *sox3^x52^* was a gift from the Bruce Riley lab.

### Whole-mount *in situ* hybridization, hybridization chain reaction and immunofluorescence

Traditional in situ hybridization (ISH) was performed as previously described (Thisse and Thisse, 2008). Following colorization, embryos were stepwise dehydrated, washed and stored in methanol. For immunofluorescence, embryos were stepwise rehydrated and blocked in 10% goat serum. Embryos were incubated overnight with anti-GFP (Aves labs, GFP-1020; 1:500), then washed and incubated with secondary antibody [goat anti-chicken-488 (Invitrogen, A-11039; 1:250)] in blocking buffer. Embryos were imaged on either a Zeiss Axioplan2 or a Axio Imager M2 with 40x objective. Deposition distances were measured in FIJI (ImageJ) and plotted in PRISM (GraphPad). In situ hybridization chain reaction (HCR) was performed with pairs of 20 probes purchased from Molecular Instruments and in accordance with their zebrafish protocol (https://files.molecularinstruments.com/MI-Protocol-RNAFISH-Zebrafish-Rev10.pdf).

Embryos were stained with Hoechst dye (1:5000), mounted in glycerol then imaged using a Leica SP5 II confocal microscope with a 20x objective (0.3 NA) and a 2.5x zoom. Immunofluorescence with rabbit anti-sox2 (Abcam, ab97959) and rabbit anti-sox3 (GeneTex, gtx132494) was performed according to Genetex IHC protocol (https://www.genetex.com/Article/Support/Index/protocols) for zebrafish whole mount embryos using goat anti-rabbit-488 (Invitrogen, A11034) secondary antibody. Imaging was performed on the Leica SP5 II with 20x objective.

### Time-lapse microscopy, image processing and quantification

For time-lapse microscopy, embryos ranging in age between 24 and 25 hpf were first anesthetized in embryo media containing 600µM MS-222 (Sigma) and then mounted in 1% low-melt agarose (NuSieve GTG), which also contained 600µM MS-222. Time-lapse confocal imaging of embryos was done using a Nikon Ti2 inverted microscope with Yokogawa CSU-W1 spinning disk confocal, Hamamatsu Orca Flash 4 version 3 camera with a 20x PlanApo Air 0.75NA objective in an incubation stage maintained at 28.5°C. *Tg(cldnb:lyn-egfp), Tg(6xtcf:lef1-miniP:d2egfp);Tg(cldnb:lyn-mscarlet), and Tg(cldnb:lyn-egfp);TgBAC(cxcr4b:h2a-mcherry)* embryos were injected with Sox1/Sox2/Sox3 morpholinos for live imaging. Sox2/Sox3 and Sox1/Sox2 morphants were generated by injecting (2.5ng sox2 MO1 and 1.0ng sox1 or sox3 MO) into single-cell embryos obtained from the aforementioned transgenic lines. 488nm and 561nm excitation lasers were used for live imaging of the pLLP and images were acquired at 5-minute time intervals over a period of approximately 14-16 hours. The sample stage was manually shifted to prevent the sample from going out of the imaging frame. Acquired images were then “Standard Deviation” projected and stitched using a custom in-house macro in Fiji before being bleach-corrected using the “Simple Ratio” technique with respect to the first timepoint. Images in the main paper were enhanced for clarity by varying their brightness and contrast without affecting their raw pixel values.

## Supporting information

video-3 Wnt activity higher mag

Video-1 CldnB-lynGFP

video-2 Wnt activity lower mag

## ACKNOWLEDGEMENTS

This work was funded by the intramural program of the *Eunice Kennedy Shriver* National Institute of Child Health and Human Development 1ZIAHD001012. Julia Rashid for help with editing. This work was initiated by Kyeong-won Yoo.

---

**Supplementary Figure 1.**
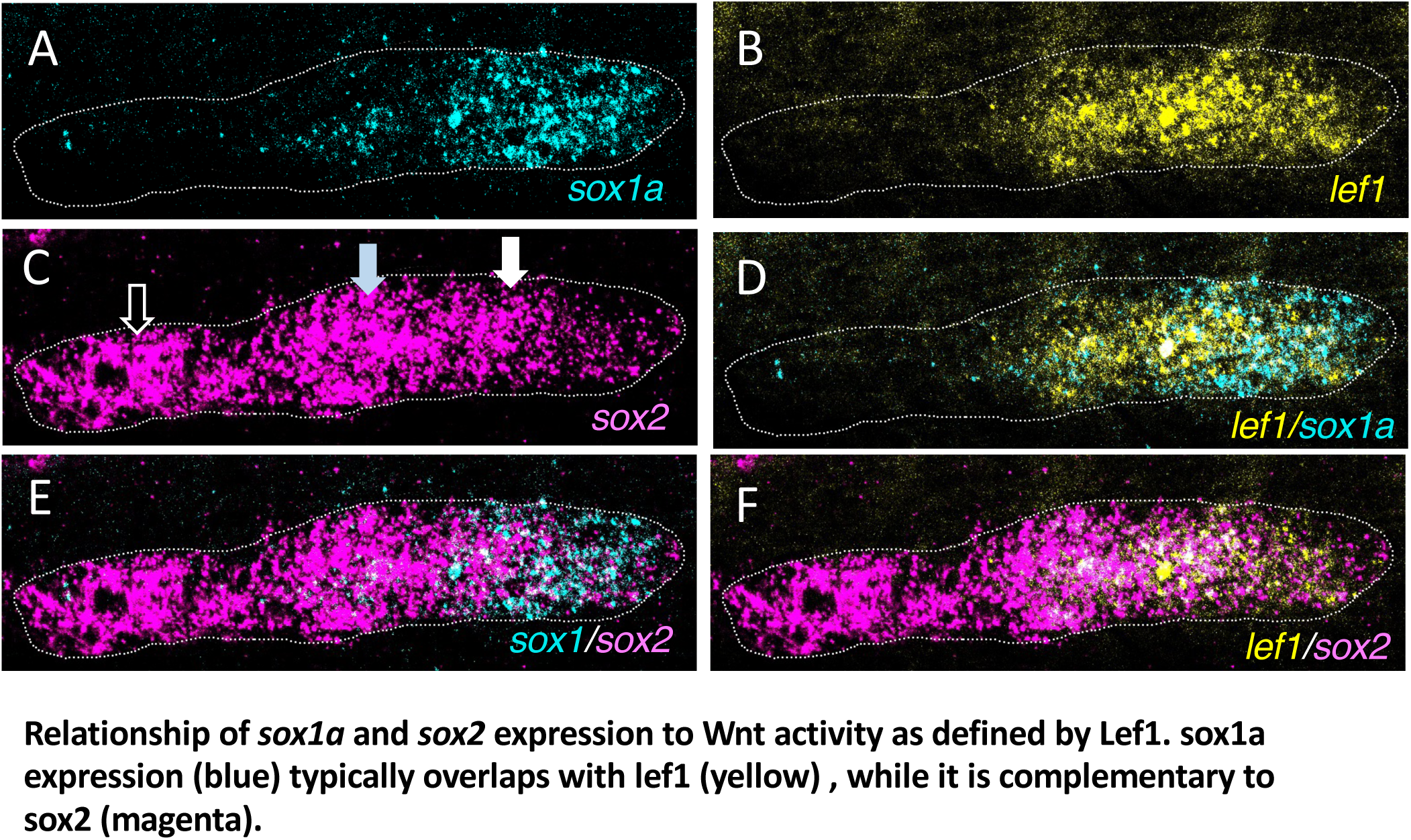
The relationship between *sox1a* and *sox2* expression and Wnt activity, as indicated by *lef1*. **A, B, D.** *sox1a* expression (cyan) typically overlaps with *lef1* expression (yellow). **A, C, E.** *sox2* expression (magenta) is complementary to *lef1* expression (yellow). **F.** *sox1a* (cyan) and *sox2* (magenta) expressions are complementary to each other. In C, white arrow shows center-biased expression, blue arrow -broad expression, and black arrow shows donut shaped expression.

**Supplementary Figure 2.**
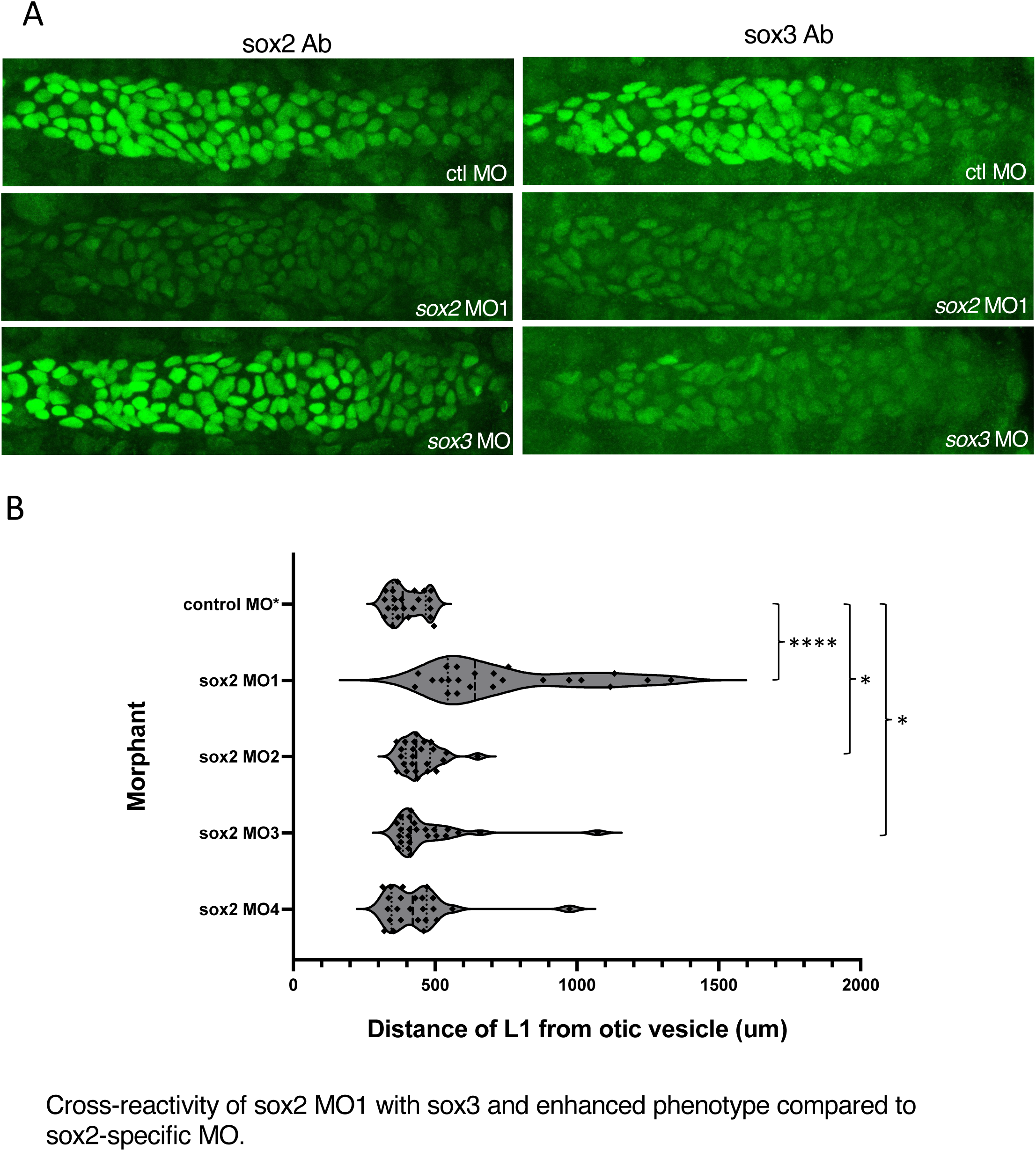
Cross-reactivity of sox2 MO1 with sox3 and enhanced phenotype compared to sox2-specific MO. **A.** sox2 and sox3 antibodies labeling control, sox2 MO1 and sox3 morphant pLLP. **B.** L1 deposition in sox2 MO1 and sox2-specific morphants. Asterisks indicate significance by Mann-Whitney test (**** p< 0.0001, * p< 0.05).

**Supplementary Figure 3.**
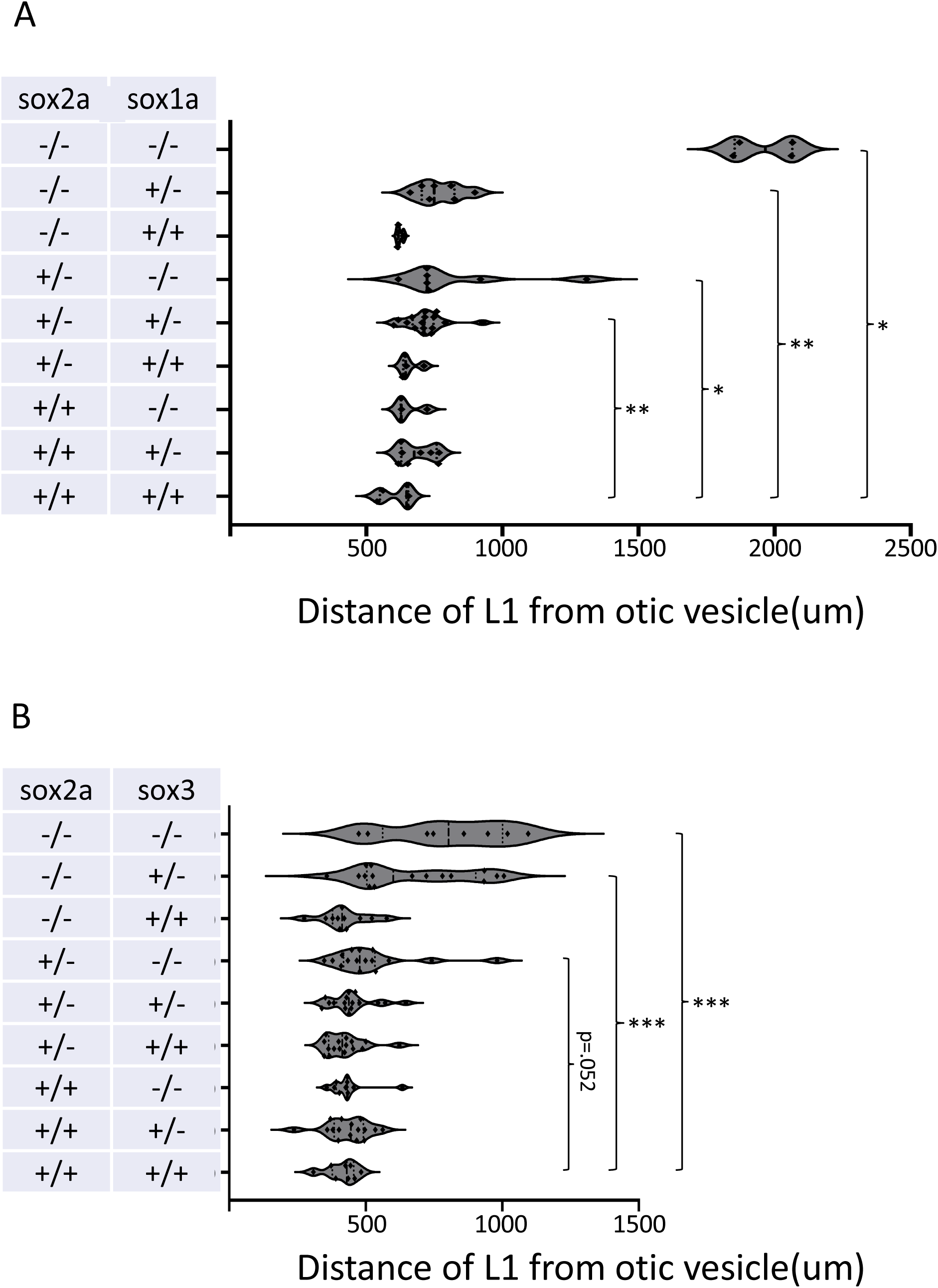
L1 deposition in sox1a/sox2 and sox2/sox3 heterozygous mutant incross. **A, B.** Deposition distance of L1 neuromast in 52hpf double mutant embryos. *sox2*^+/-^ and *sox1a*^+/-^ or *sox3*^+/-^ were crossed to generate heterozygous progeny for imaging followed by genotyping. Asterisks indicate significance by Mann-Whitney test (*** p< 0.001, ** p< 0.01, * p< 0.05).

**Supplementary Figure 4.**
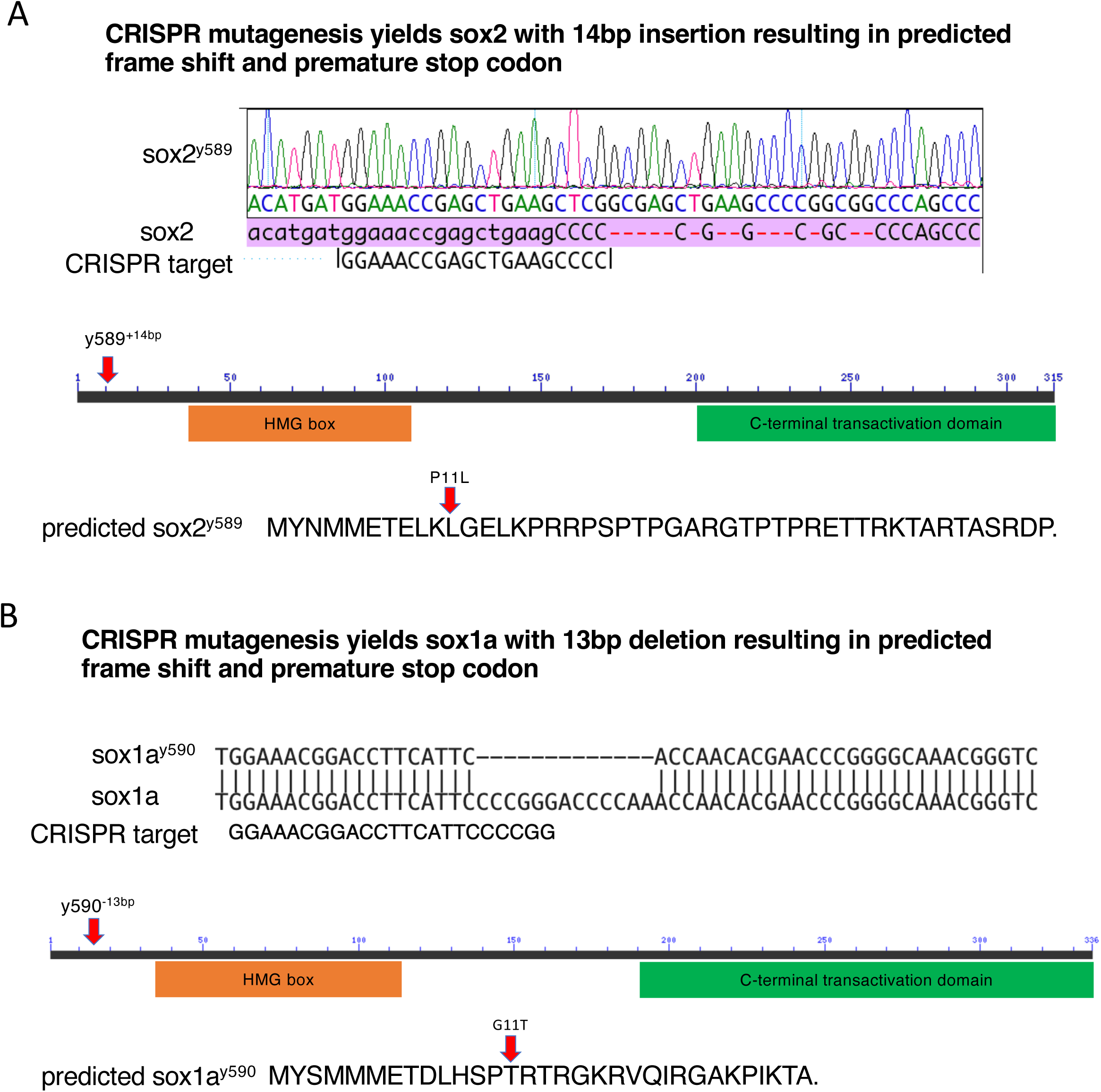
CRISPR mutagenesis of sox2^y589^ and sox1a^y590^. **A.** sox2 insertional mutant with site of frame shift and predicted translation. B. sox1a deletional mutant with site of frame shift and predicted translation.

## A Supplementary Videos

**Supp. Video 1**.

**Membrane (Ctl + Sox2/3) video** Timelapse confocal imaging of representative Control MO-injected (Top) and Sox2 MO1/Sox3 MO-injected (Bottom) embryos showing differences in pLLp migration and neuromast deposition. Imaging was done from ∼24 hpf to ∼42 hpf. Regions of high intensities are shown in warm colours.

**Supp. Video 2**

**Membrane + Lef1 reporter entire timelapse video:** Timelapse confocal imaging of representative Control MO-injected (Top) and Sox2 MO1/Sox3 MO-injected (Bottom) *Tg(6xtcf:lef1-miniP:d2egfp);Tg(cldnb:lyn-mscarlet)* embryos showing differences in Lef1 reporter activity in the pLLp, with regions of high intensities of Lef1 reporter shown in warm colours, and membrane shown in grey. Imaging was done from ∼24 hpf to ∼40 hpf.

**Membrane + Lef1 reporter focused primordium video (Supp. Video 3):** A closer look at the embryos (Control MO-injected (Left), and Sox2 MO1/Sox3 MO-injected (Right)) in Supp. Video 2 focusing on the pLLp and deposited neuromasts. Neuromasts that are deposited first are circled blue, while those that are deposited next are circled red.

